# Significant reduction in blackfly bites following implementation of Slash and Clear: An option to consider for onchocerciasis elimination in areas of persistent transmission

**DOI:** 10.1101/2023.03.27.534316

**Authors:** André Domche, Hugues C. Nana Djeunga, Philippe B. Nwane, Guy R. Njitchouang, Betrand Nono Fesuh, Flobert Njiokou, Benjamin Jacob, Sébastien D. Pion, Joseph Kamgno

## Abstract

**Background:** Although “Slash and Clear” has proven effective in reducing blackfly densities in low transmission foci, the impact of this strategy in high transmission settings with large rivers and important vector densities remains to be demonstrated.

**Methods/Principal findings:** A controlled before-and-after community-based intervention comprising two arms (Bayomen as control site and Biatsota as intervention site) was carried out in the Mbam Valley (Centre Region, Cameroon). In each arm, baseline blackfly densities were collected over one year using the human landing method. The intervention consisted of destroying the trailing vegetation where blackflies breed. Blackfly densities were collected post-intervention to assess the impact of the intervention. Before the intervention, a total of 36,273 and 29,041 blackflies were collected in Bayomen and Biatsota, respectively. After the intervention period, the total blackfly density in the intervention site decreased from 29,041 to 20,011 (31.1% reduction), while an increase of 2·7% was observed in the control site (from 36,273 to 37,248). The Poisson mixed regression model shows that the reduction was significantly greater in the intervention site than in the control site (p<0.0005).

**Conclusions/Significance:** This study showed that “Slash and Clear” approach is feasible and has a significant impact on vector densities in a high transmission setting. Further studies are needed to investigate the long-term impact of this vector control approach, and how this promising strategy can be scaled-up and sustained until elimination of onchocerciasis.

**Author summary:** River blindness persists in some foci in Cameroon despite more than two decades of ivermectin-based preventive chemotherapy. Mass drug administration (MDA) appears insufficient to interrupt onchocerciasis transmission in these hotspots, and should be complemented by vector control, the most promising alternative strategy to date. In 2018, the effectiveness of a new community-based vector control approach, known as slash and clear, was demonstrated. This strategy involves the removal of trailing vegetation at breeding sites, a primary attachment points for blackfly larvae. In this study, we show that this environment-friendly intervention is feasible and has a significant impact on blackfly densities in high transmission settings. This promising intervention can be combined with regular annual ivermectin-based preventive chemotherapy to accelerate onchocerciasis elimination.

## Introduction

Onchocerciasis, better known as river blindness, is a parasitic disease caused by a nematode called *Onchocerca volvulus*, which is transmitted to humans by black flies of the genus *Simulium* [1]. These blackflies breed in fast-flowing streams and rivers, usually near remote rural villages. Onchocerciasis infection can cause skin disease, including severe itching, rashes, or nodules under the skin, as well as visual impairment and, lately irreversible blindness [2]. Because of these clinical manifestations and the associated socio-economic burden, onchocerciasis is considered a public health problem in endemic countries. The disease is endemic in 31 countries in sub-Saharan Africa, South America, and Yemen[3, 4]. The World Health Organization (WHO) estimates that 220 million people worldwide are at risk of onchocerciasis and therefore require preventive chemotherapy, with 99% of those infected living in Africa. An estimated 14·6 million of those infected suffer from skin disease, and 1·15 million have vision loss [5].

Onchocerciasis control currently relies on targeting the causative agent through a strategy known as Community-Directed Treatment with Ivermectin (CDTI) [6]. WHO recommends annual distribution of ivermectin for at least 15-17 years with high coverage (≥80%) to expect interruption of transmission, as evidence by very low incidence of the infection in children aged 5-9 years [7]. Based on this strategy, onchocerciasis has been eliminated in several foci in Africa[8–12]. However, despite more than 20 rounds of uninterrupted annual CDTI in the Mbam valley (of Cameroon), the disease persists with microfilarodermia prevalence up to 50% in some villages [13]. This persistence of infection could jeopardize the WHO’s goal of verifying the interruption of onchocerciasis transmission in 31% of endemic countries by 2030 [14].

Alternative/complementary strategies, including vector control, have been recommended to accelerate onchocerciasis elimination in these hotspot areas in a timely manner [15]. Recently, a low-cost, community-based, and environment-friendly approach (Slash and Clear) to tackle *O. volvulus* vector was shown to be effective by drastically reducing blackfly densities and the resulting transmission of infection [16]. However, this trial was conducted in a hypo-endemic area of onchocerciasis in northern Uganda, where the vector breeds in small rivers and their breeding sites easily accessible. This raised interest in the feasibility and impact of such a community-based intervention in areas with a more important hydrographic network of rivers with very large sections.

This study aims to assess the impact of the newly designed community-based vector control (Slash and Clear) on blackfly densities in a high onchocerciasis transmission area in the Mbam Valley, Centre Region, Cameroon.

## Materials and methods

### Study area

The present study was conducted in the Bafia Health District, located in the Mbam Valley, Centre Region, Cameroon. It is a transition zone between forest and savannah, irrigated by many fast-flowing rivers, mainly the Sanaga and its main tributary, the Mbam (Fig. 1). The total length of the Mbam river is 494 km, and its course is interspersed by many rapids and falls, which are suitable for breeding sites of black flies [17]. The climate is equatorial with a long dry season from November to February, a short rainy season from March to June, a short dry season from June to August and a long rainy season from August to November. The vectors of *O. volvulus* in the Bafia Health District are the forest-dwelling species *Simulium squamosum* and *Simulium mengense* [18].

**Fig 1.**
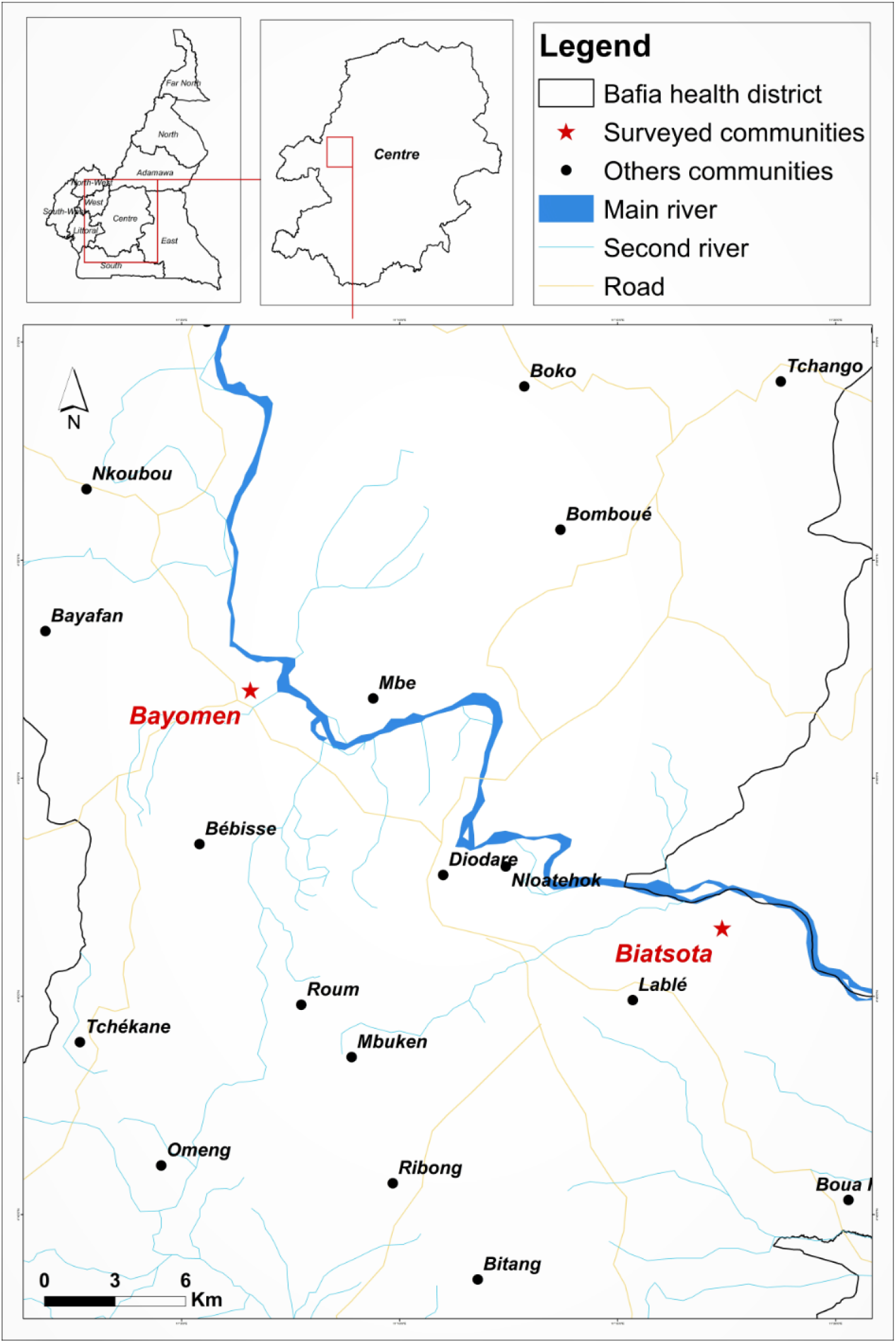
Study area with surveyed communities

According to the National Onchocerciasis Control Programme (NOCP), CDTI was initiated in Bafia Health District in 2000 and has been maintained at an annual frequency since then [19]. Surveys conducted in 2015 and 2019, after 15-19 rounds of annual uninterrupted CDTI, revealed microfilarial prevalence ranging from 24.4 to 57.0% and 10.7-54.2% in the surveyed villages [13] (Kamgno et al., unpublished data).

### Study design

This study was a controlled before-and-after community-based intervention with two arms: an intervention arm (Slash and Clear implemented in the village of Biatsota), and a control arm (no Slash and Clear in the village Bayomen). Prior to the intervention, a baseline collection of black flies was performed over one year to assess the natural population dynamics and better evaluate the impact of the intervention.

### Baseline blackfly collection

To establish a baseline of the vector population density, blackfly collections were conducted at a monthly frequency over one year, from October 2019 to December 2020, prior the introduction of “Slash and Clear”. Collections were conducted using the human landing collection method [20], by a team of two operators per collection site working alternately between 07:00 and 12:00 hours for the first, and between 12:00 and 17:00 hours for the second. Female blackflies landing on collectors’ exposed legs were caught using a mouth aspirator before they had the opportunity to take their blood meal as described elsewhere [18]. Blackflies collected during each time slot were taken to the field laboratory by the field supervisor, where they were counted and identified by an experienced entomologist.

### Prospection and characterization of blackfly breeding sites

Ground and boat prospections were carried out by experienced entomologists and selected community members with good knowledge of the river and its tributaries. The aim of these prospections was to identify blackfly breeding sites and to describe their environment to characterise the type of vegetation and rocks used by female blackflies as supports for egg laying. The identification of breeding sites was guided by the conditions favourable to the establishment of blackfly larvae in a stream, including but not limited to (i) the presence of submerged substrates, (ii) the presence of a satisfactory water velocity (0.5 to 2 m/s), (iii) the presence of food particles (larvae feed passively, capturing any particles carried by the current) [21]. The prospection was carried out over two days in December 2020, approximately 2 km upstream and downstream of the village targeted for the complementary intervention (slash and Clear) [16]. All potential breeding sites (with or without larvae/pupae) were geo-referenced using a Global Positioning System [GPS eTrex; Garmin (Europe) Ltd, Southampton, UK] and characterised according to the type/nature of the laying support and the number of larvae/pupae colonising a breeding site, following the method previously used by the Onchocerciasis Control Programme (OCP) in West Africa [21, 22].

### Implementation and follow-up of “Slash and Clear”

Following prospections of breeding sites in the intervention village, and prior to the Slash and Clear exercise, the research team met with the community leaders, including the village chief and the community members who were trained to identify breeding sites, to define the profile of individuals who will carry out the “Slash and Clear”, so-called “Slashers”. A total of eight men with proven water experience, mostly fishermen, aged between 32 and 45 years were recruited. The intervention (Slash and Clear) was carried out on four consecutive days per month from December 2020 to August 2022. Equipped with their life jackets and their machetes, the Slashers cut down the trailing vegetation of the breeding sites and their surroundings, using boats when the latter were hard to reach on foot. The trailing vegetation cut was discarded on the banks of the river where any larvae/pupae present died of desiccation. When the vegetation was too heavy to be transported to the riverbanks in the boat, it was drained by the water flow to the riverbed where it was drowned with the larvae/pupae. To assess the impact of “Slash and Clear”, black fly collections were carried out monthly in the two study sites as described above, just before the next exercise.

### Statistical analyses

Statistical analyses were performed using the R software (version 4.0.2). Follow-up plots were used to monitor the evolution of the number of blackflies collected with the date of collection for both the control site (Bayomen village) and the intervention site (Biatsosa village). Poisson Mixed Regression Models with random slope for the date variable and an autoregressive homogeneous variance-covariance matrix order-1 were used to account for variations in the number of blackflies collected (dependent variable recorded as counts) in the time series. The models controlled for independent variables such as collection date and Village as fixed effects, with village playing a dual role as intervention variable (Bayomen as control and Biatsosa as intervention site) and cluster variable. Since a baseline study was initially conducted without implementing the slash and clear method, separate models were run for the baseline and intervention studies to account for confounding. No significant overdispersions was observed in the models based on overdispersion tests. The R package “glmmTMB”[23] was used to run the models. The significance threshold was set at 5% for all statistical analyses.

### Ethics statement

This study protocol was approved by the Regional Ethics Committee for Research in Human Health for the Centre Region (no. 905/CRERSHC/2019) and an administrative authorization was granted by the Centre Regional Delegation for Public Health (no. 624/AP/MINSANTE/SG/DRSPC/CRERSH). The objectives and schedule of the study were explained to the District Medical Officer, the community leaders, and all volunteers before requesting a verbal consent to participate was obtained. Participation was entirely voluntary, and all individuals were free to opt out without fear of retaliation from their community leaders or programme staff.

## Results

### Dynamics of blackfly densities in the two villages prior to the intervention

Before to the intervention, a total of 65,314 (median: 2,101; Interquartile range, IQR: 1,206-2,535) blackflies were collected in the two study sites over one year (October 2019-December 2020). In Bayomen, 36,273 (median: 2,235; IQR: 1,426-2,544) blackflies were collected, and in Biatsota 29,041 (median: 1,912; IQR: 1,206-2,203) blackflies were collected. Black flies were present throughout the year in both villages, with varying densities from one month to another, the highest densities being recorded in both villages at the beginning of the dry season (in November). The monthly dynamics of black flies followed a similar trend in both villages (Fig. 2).

**Fig 2.**
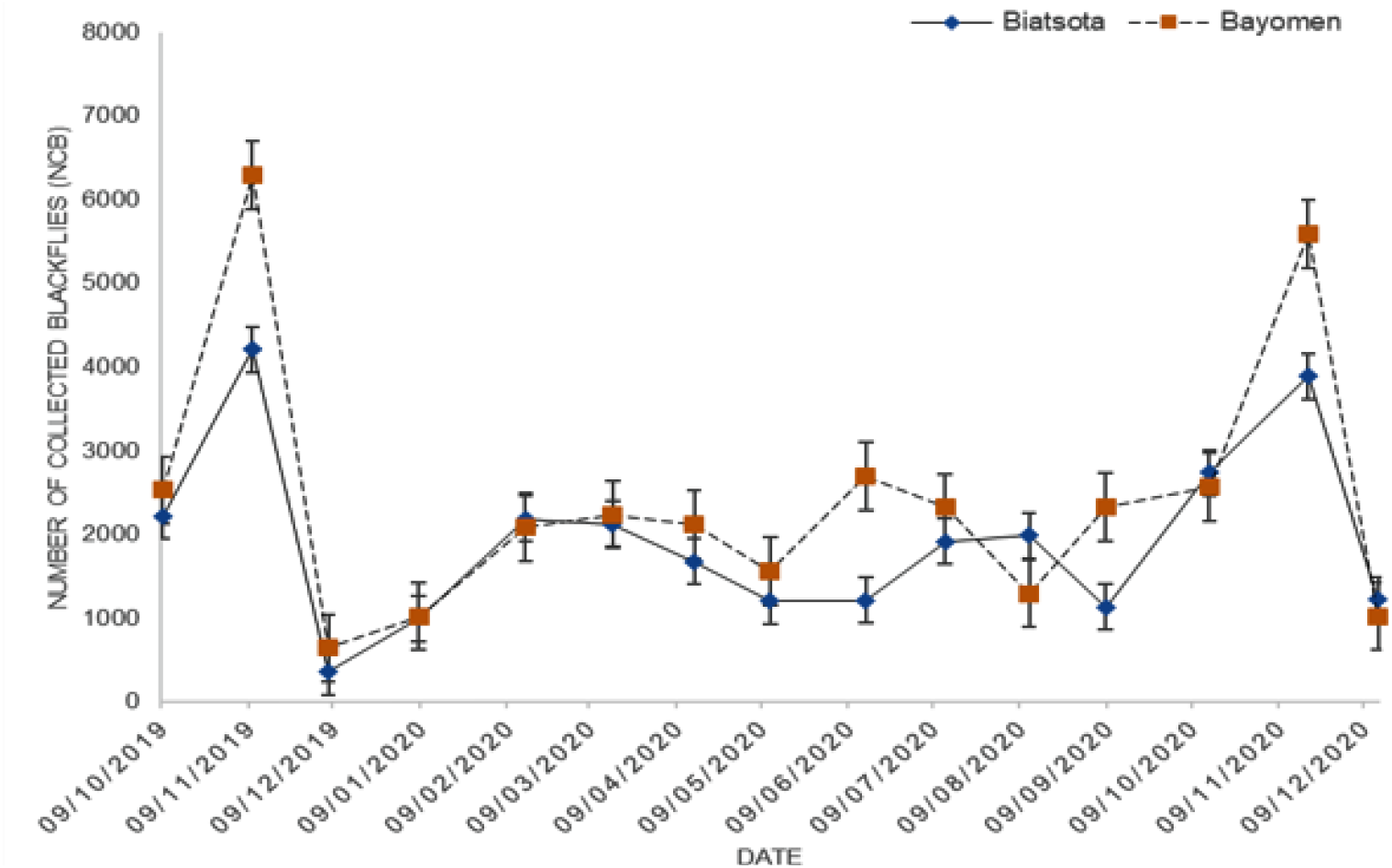
Dynamic of blackflies before intervention in the two villages

### Blackfly breeding sites identification in the intervention area

A total of seven breeding sites were identified in the Biatsota village, approximately 2 km upstream of the Mbam River, while none were identified downstream. Five of these breeding sites were heavily colonised (≥ 50 larvae/pupae) and one was lightly colonised (≤ 10 larvae/pupae) (Fig. 3). The last and largest potential breeding site was inaccessible due to the volume of water and velocity of the stream.

**Fig 3.**
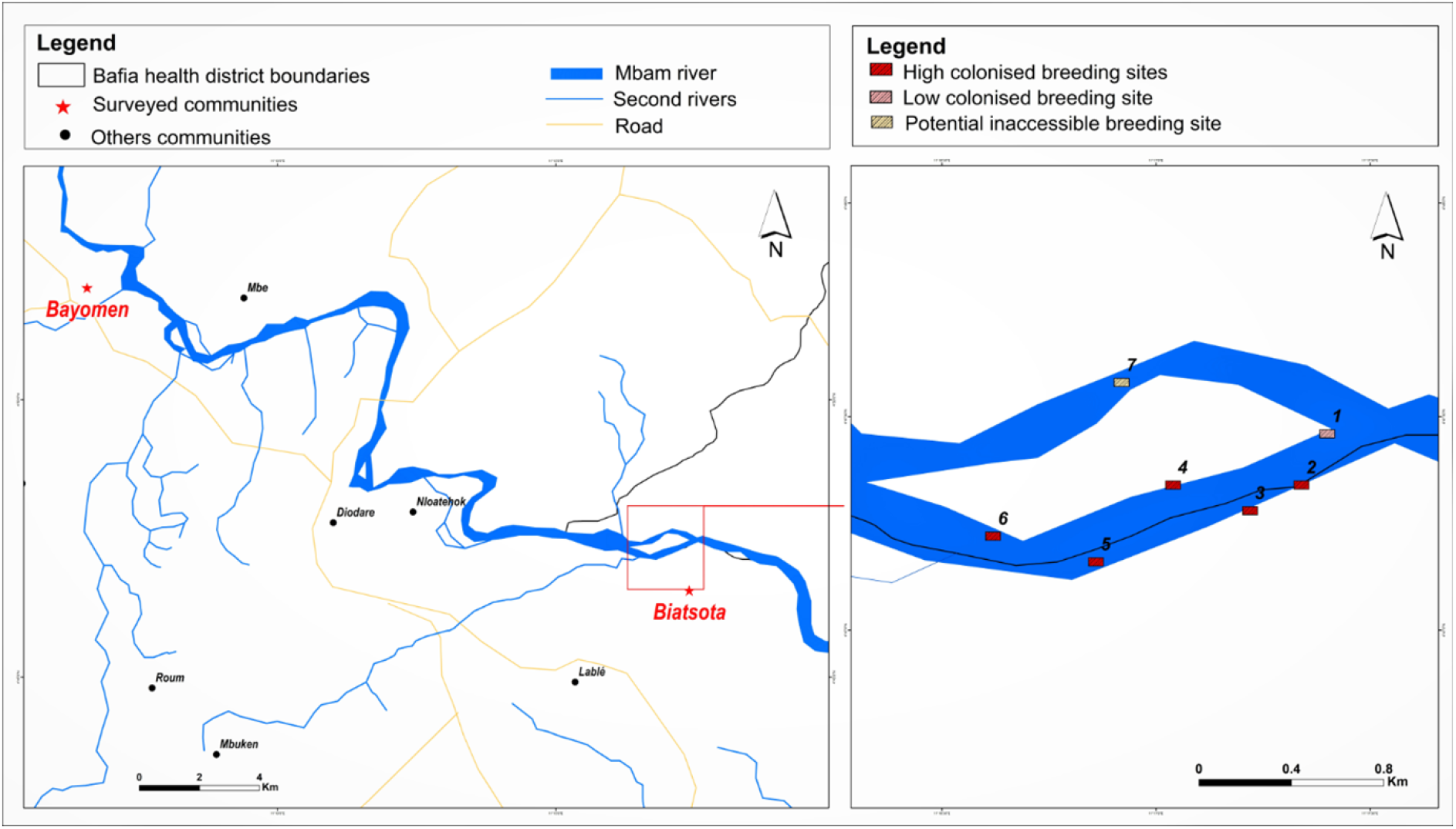
Distribution of potential identified breeding sites.

The main supports for blackfly oviposition were plants, particularly *Pandanus candelabrum*, which sheltered more than 90% of the larvae/pupae (Fig. 4).

**Fig 4.**
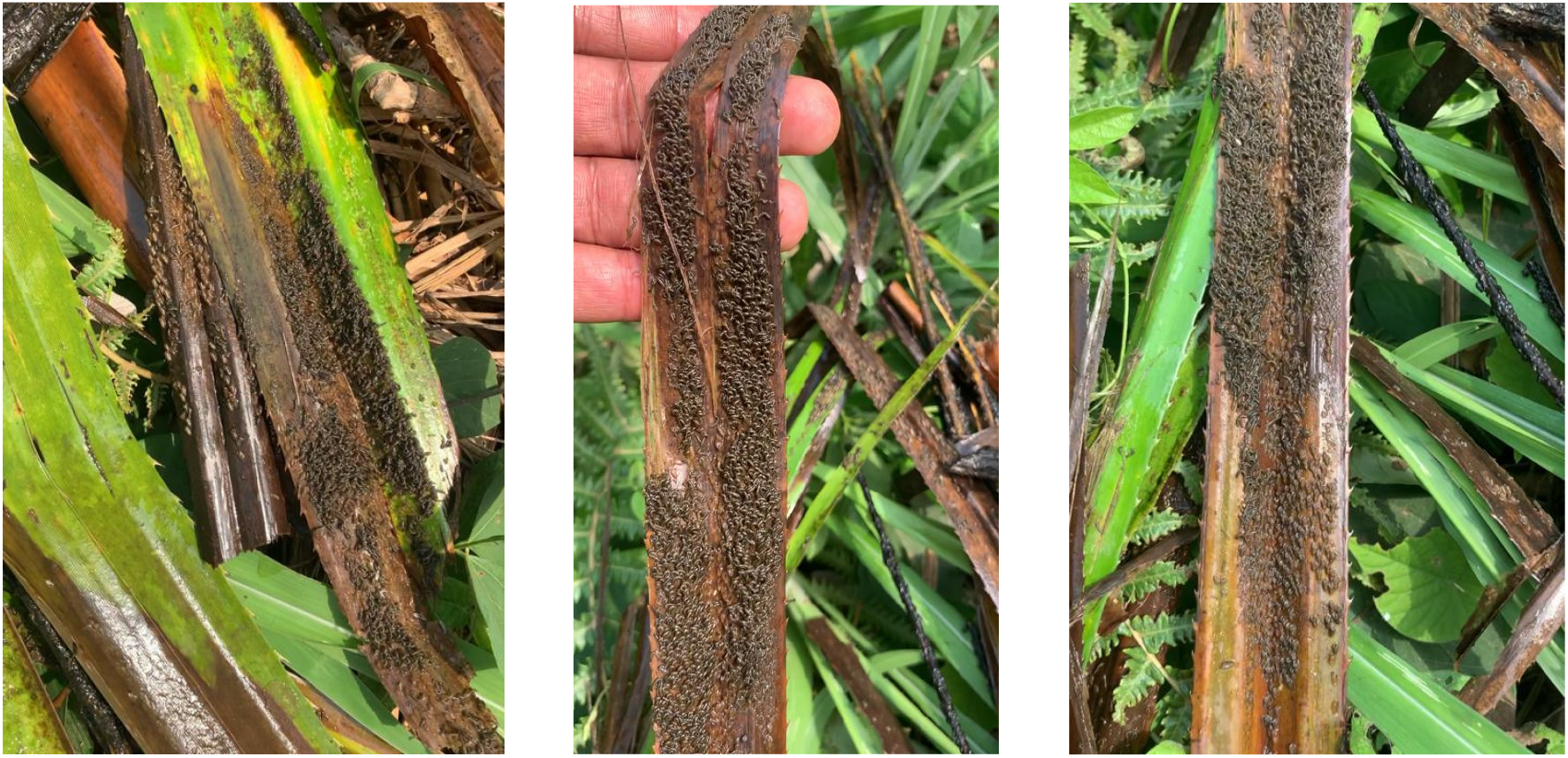
*Pandanus candelabrum* with black fly larvae/pupae

### Dynamics of blackfly densities in the two villages during the intervention

During the intervention (January 2021-November 2021), a total of 57,259 (median: 1,068; IQR: 798-1,776) blackflies were collected in the two villages; 37,248 (median: 1,659; IQR: 1,043-2,060) in Bayomen and 20,011 (median: 890; IQR: 692-1,072) in Biatsota. Figure 4 illustrates the overall dynamics of blackfly densities during the baseline and intervention phases in both control and intervention sites (Fig. 5).

**Fig 5.**
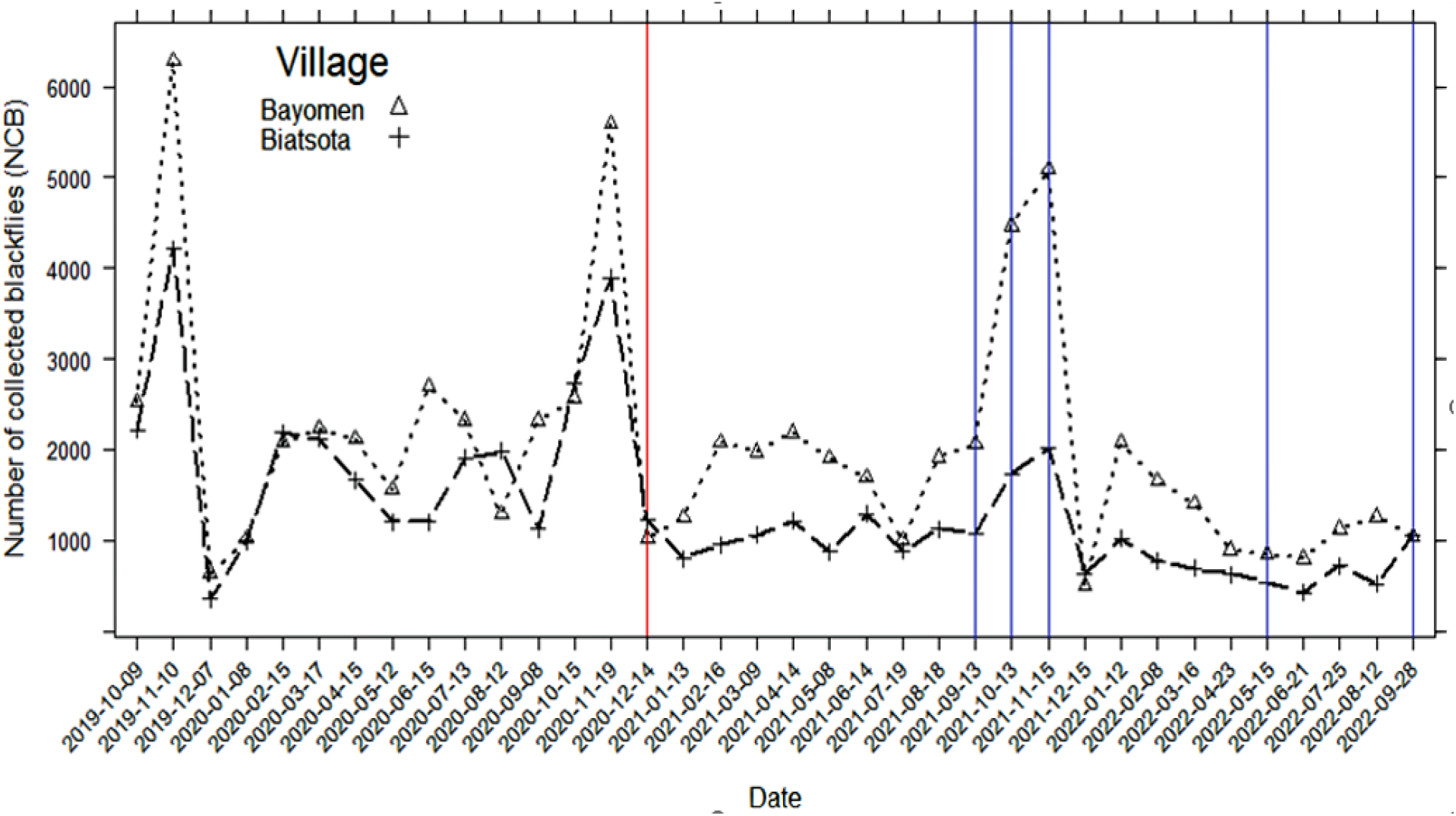
Dynamics of monthly blackfly collection during baseline and intervention phases in both control (Bayomen) and intervention (Biatsota) sites. Red line represents the beginning of the intervention and purple lines are the months without intervention.

While blackfly densities followed the same trend and somehow overlapped in both sites during the baseline phase, the Poisson mixed model (Table 1) showed that fly densities were significantly lower in the intervention site (Biatsota) compared to the control site (Bayomen) (*p*<0.0005). In fact, black fly densities were reduced by 31.1% in Biatsota while an increase of 2·87% was observed in Bayomen. The intervention didn’t take place during the months of September, October, and November 2021 due to the increased volume of water with the onset of the rains. Nevertheless, the number of flies collected in the intervention village during this period remained low compared to the control village (4,826 vs 11,614 black flies) and the preintervention period in the same village (4,826 vs 7,756 black flies).

**Table 1.**
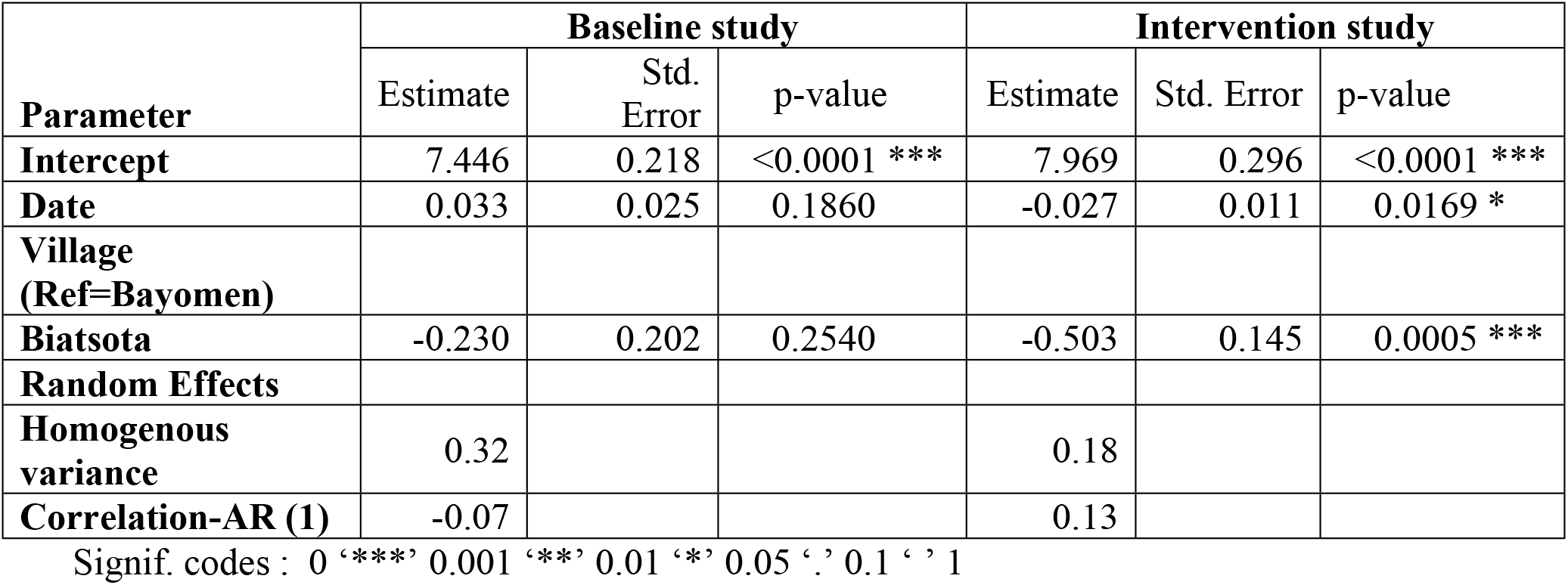
Effect of intervention on the number of collected blackflies in the control and intervention villages, controlling for date of collection and baseline study. Poisson Mixed Regression Models with random slope for the date variable with autoregressive order-1 homogeneous variance-covariance matrix

## Discussion

The aim of this study was to evaluate the impact of community slash and clear on blackfly densities in a high onchocerciasis transmission area in central Cameroon.

The baseline data recorded in this study revealed very high biting rates in Bafia Health District as previously described [18], and compared to biting rates recorded in other similar settings (first-line communities with no vector control) [24, 25]. The Mbam river appears to be one of the major sources of onchocerciasis vectors in the country. These high biting rates reflect a high level of blackfly nuisance to the human population as was confirmed by the communities themselves during a study conducted in this area in 2019 [26] and may contribute significantly to the persistence of the disease in this focus [13]. In fact, a quantitative study of community knowledge/perceptions and attitudes/practices regarding onchocerciasis and blackfly nuisance and bioecology revealed that, in addition to the transmission of the *O. volvulus* parasite, blackflies are responsible for a significant nuisance that reduces population productivity [26]. In addition, a significant proportion of people surveyed indicated that they would be willing to help control blackflies. This suggests both a need and an opportunity to implement a community-based vector control strategy in such areas.

The monthly variability of blackfly densities confirms the highly dynamic nature of the population and can be explained by the water discharge in the river, which are strongly correlated with climatic conditions, mainly precipitation and temperature [27]. In fact, at the end of the rainy season, the volume of water decreases, and the egg-laying supports become more accessible to female blackflies, that can easily and safely lay their eggs. Under such conditions, most of the eggs will reach the end of their development in about 9 days, thus increasing the density of emerging adults. As the control and intervention sites are approximately 25 km apart, they are likely experience to the same climatic conditions and the same variation in the volume and velocity of the Mbam river, which explains the similar trend observed in the monthly dynamics of blackfly densities.

It is well known that the female of *S. damnosum sensu lato* species complex lays her eggs on rocks or vegetation which develop into larvae in fast-flowing rivers and streams [28]. Since the downstream of the Mbam river is low flowing, and there are no rocks or other supports to increase the speed of the water, no breeding sites were found 2 km downstream of the river during the prospections. The large number of breeding sites observed 2 km upstream of the river is explained by the presence of trailing vegetation and rocks, as well as the average water discharge of 7,100m^2^/s which generates oxygen necessary for the development of immature stages [17].

The reasons why *Pandanus candelabrum* was the preferred attachment site for immature blackfly stages are yet to be established. However, it can be hypothesized that the physical structure and/or the composition of this plant, which is likely to be suitable for the development of immature blackfly stages, or the fact that it is the most abundant plant in areas may be the reasons why it was chosen by adult female blackflies to lay their eggs. Further research into the chemical composition of this plant may provide a better explanation for this phenomenon. This predominance of larvae on this plant suggests that its physical removal along the watercourse, especially in an environment where all the conditions for larval development are met, may deprive the blackflies of their preferred attachment sites. This, in turn, should drastically reduce the population of the vector that successfully breeds to maturity. The heavy larval/pupae colonization of most breeding sites (5/6 accessible) reflects the high blackfly densities observed in the study communities.

The current study showed that removing the trailing vegetation that serves as a substrate for immature blackflies reduced the median density of adult vectors by 53.5%. Although significant, this reduction was less important than the 89-99% reduction in blackfly numbers reported in northern Uganda [16]. In fact, Slash and Clear activities were carried out 1km upstream and downstream of a target village in Uganda, and the rivers where the intervention was carried out were small, with a small number of breeding sites all accessible, whereas the Mbam river is very large, with many breeding sites, some being inaccessible. Despite the impact of the intervention on blackfly densities, an important proportion (68.9%) of blackflies survived to the intervention. Although it can be hypothesized that not every single breeding site was not destroyed, the repopulation may come from the upstream neighbourhood (more than 2 km away) where the intervention was not carried out. These observations highlight the difficulties to implement this strategy in large rivers and should be considered when designing the scale-up and sustainability plans for such a vector control approach.

Three months after the intervention was stopped (from September to November 2021) due to the increase in water volume, the number of flies collected in the intervention village remained low compared to the control village. This shows that the impact of the intervention might last longer than 2-3 months and raises some questions about the frequency of the intervention (once a month in this study), which could be reduced to once every two or three months to facilitate the implementation of the intervention, especially for the scale-up and sustainability perspectives.

### Lessons learned for scale-up and sustainability

Although difficult to implement in large rivers due to the high water discharge and the inaccessibility of some breeding sites, several potential tricks can be proposed to minimize the impact of these obstacles, including but not limited to (i) maximising Slash and Clear activities during the dry season, when water levels and discharge are sufficiently low; ii) taking into account the activities of hydroelectric dams located upstream of the target villages while implementing Slash and Clear; iii) using remote sensing imagery prior to ground/boat prospection to maximize the identification of areas with characteristic features of S. *damnosum* s.l. breeding habitat [29]; iv) integrating targeted ground larviciding component using drones and biological compounds such as *Bacillus thuringiensiss israelensis*, for inaccessible sites whatever the season; v) engaging the community and local government or council to ensure scale-up and sustainability; vi) using better equipment for navigation and clearing of the breeding sites. One of the key assets of this strategy is the fact that the community has agreed to participate in blackfly control [26], provided that they are equipped and trained.

## Conclusions

This study demonstrated that this community-based environment-friendly vector control approach (Slash and Clear) is largely feasible in a high transmission setting and has a significant impact on vector densities. Further studies are needed to investigate the long-term impact of this vector control strategy, and how best to minimize the various challenges encountered to optimally complement mass drug administration and therefore accelerate onchocerciasis elimination in high transmission settings.

## Acknowledgements

The authors would like to thank the German Federal Ministry for Economic Cooperation and Development (BMZ) through KfW for supporting this study, and the *Organisation de Coordination pour la lutte contre les Endémies en Afrique Centrale* (OCEAC) for managing the study in Cameroon.

## Author Contributions

AD, HCND, and JK designed the study protocol. AD, HCND, BNF, SP and JK contributed to the drafting and revision of the manuscript. AD, HCND and BNF prepared the first draft of the manuscript and conducted data analysis. AD, HCND, GRN, BJ and PBN monitored data collection and management. All the authors critically reviewed the results and contributed to the interpretation of data. All authors participated in the interpretation of results and revisions of the manuscript. All authors read and approved the final manuscript.

